## Supplemental text and figures for "Fast Objective Coupled Planar Illumination Microscopy"

### Aberrations for non-focal imaging

November 21, 2018

Consider an optical system that satisfies the Abbe sine condition for two planes, called the object focal plane and image focal plane. Consider a point source along the optic axis separated by $\Delta z$ from the object focal plane. Rays are emitted from this object over a range of angles and can be traced to the object focal plane.

The sine condition states that each ray satisfies

$$n \sin \theta = M n' \sin \theta', \quad (1)$$

where $n$ is the refractive index of the object immersion medium, $n'$ that of the imaging immersion medium (here taken to be 1 for air), $M$ the local linear magnification near the axis, and $\theta$, $\theta'$ are the ray angles in the object and image space, respectively. For a ray of angle $\theta$, let $h$ denote the height of the strike position of each ray in the object focal plane, where $h = \Delta z \tan \theta$. We can

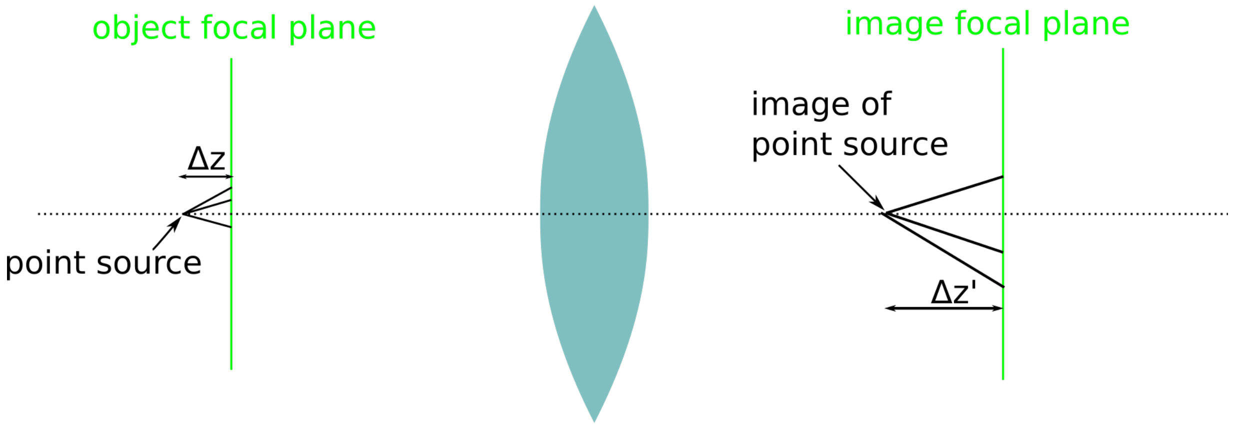

calculate the position $\Delta z'_\theta$ where this intersects the optic axis:

$$\begin{aligned}
\Delta z'_\theta &= \frac{h'}{\tan\theta'} && (2)\\
&= \frac{Mh}{\frac{\sin\theta'}{\sqrt{1-\sin^2\theta'}}} && (3)\\
&= \frac{M\Delta z\tan\theta}{\frac{\frac{1}{M}n\sin\theta}{\sqrt{1-\frac{1}{M^2}n^2\sin^2\theta}}} && (4)\\
&= \frac{M^2}{n}\Delta z\frac{\sqrt{1-\frac{1}{M^2}n^2\sin^2\theta}}{\sqrt{1-\sin^2\theta}}. && (5)
\end{aligned}$$

Note that the strike position is a function of $\theta$, and therefore in general we do not have perfect focus. The only time the angle-dependence disappears is when the ratio of square-roots cancels, which requires $|M| = n$. This is the condition derived more generally by the Maxwell perfect-imaging theorem. (We can also see that the longitudinal magnification is $\frac{M^2}{n}$.)

When this condition does not hold, the fractional range of strike positions (up to the maximum angle consistent with the numerical aperture NA) is

$$\frac{\sqrt{1-\frac{\mathrm{NA}^2}{M^2}}}{\sqrt{1-\frac{\mathrm{NA}^2}{n^2}}} - 1 \qquad (6)$$

For $M = 20$ and water immersion, at NA 0.5 we get approximately 8%, and at NA 1 we get a 50% spread.

The axial RMS spot radius offers a more informative summary of the imaging performance. This quantity is the square root of the mean squared error (MSE) in strike position for every ray within the collection cone of the objective lens. For a ray with angle $\theta$ emitted from a source at $\Delta z$ the squared error is

$$(\Delta z'_\theta - z'_0)^2 = \left(\Delta z'_\theta - \frac{M^2}{n}\Delta z\right)^2 \qquad (7)$$

For each angle $\theta$ an ideal point source at $\Delta z$ emits a circle of rays. It can easily be shown with trigonometry that the total light collected at angle $\theta$ is proportional to $\tan(\theta)$. Therefore we scale the error at each angle $\theta$ accordingly to get the sum squared error and then divide to calculate MSE:

$$\begin{aligned}
\mathrm{MSE} &= \frac{\int_0^{\theta'}\tan(\theta)\left(\Delta z'_\theta - \frac{M^2}{n}\Delta z\right)^2 \mathrm{d}\theta}{\int_0^{\theta'}\tan(\theta)\,\mathrm{d}\theta} && (8)\\
&= \frac{\int_0^{\theta'}\tan(\theta)\left(\Delta z'_\theta - \frac{M^2}{n}\Delta z\right)^2 \mathrm{d}\theta}{-\ln(\cos(\theta'))} && (9)
\end{aligned}$$

The upper limit, $\theta'$ is the maximum angle that the objective can collect as specified by the NA. Note that the above calculates the RMS spot radius in image space; in order to convert this to object space units divide by the axial magnification.

Fig S1

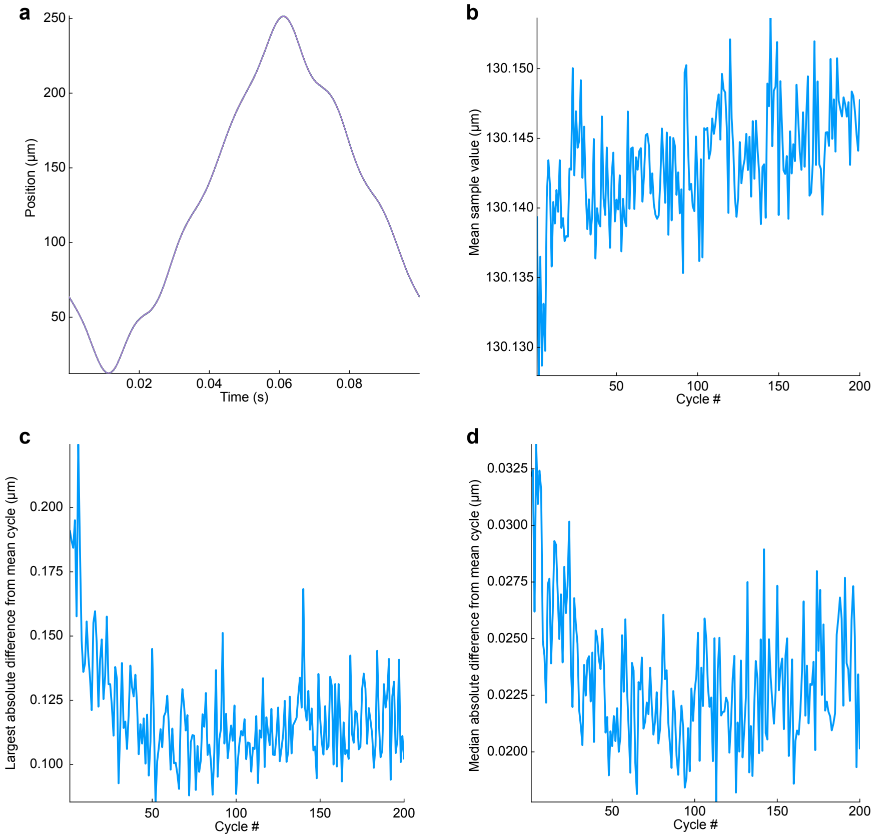


### Fig. S1

Consistency of the piezo response

**(a)** Overlay of 200 consecutive piezo scan cycles as measured by the capacitive sensor of the device (cycles during the first 20 s of operation are excluded). The command signal was a 10 Hz triangle wave. Samples were acquired at 100 kHz and downsampled to 10 kHz for plotting.

**(b)** Closed-loop control of the piezo prevents drift ("creep") in the mean piezo response measured during each cycle from (a).

**(c)** Piezo response cycles are also consistent on a per-sample basis after an initial settling period. First a mean response cycle was created by averaging corresponding samples across the 200 cycles. Plotted is the maximum absolute difference of any sampled value of each cycle from the corresponding value in the mean cycle. Before computing differences each cycle was lowpass filtered with a gaussian kernel of width 100 µs to reduce sampling noise.

**(d)** Similar to (c), but the median difference between corresponding samples in each cycle is shown.

Fig. S2

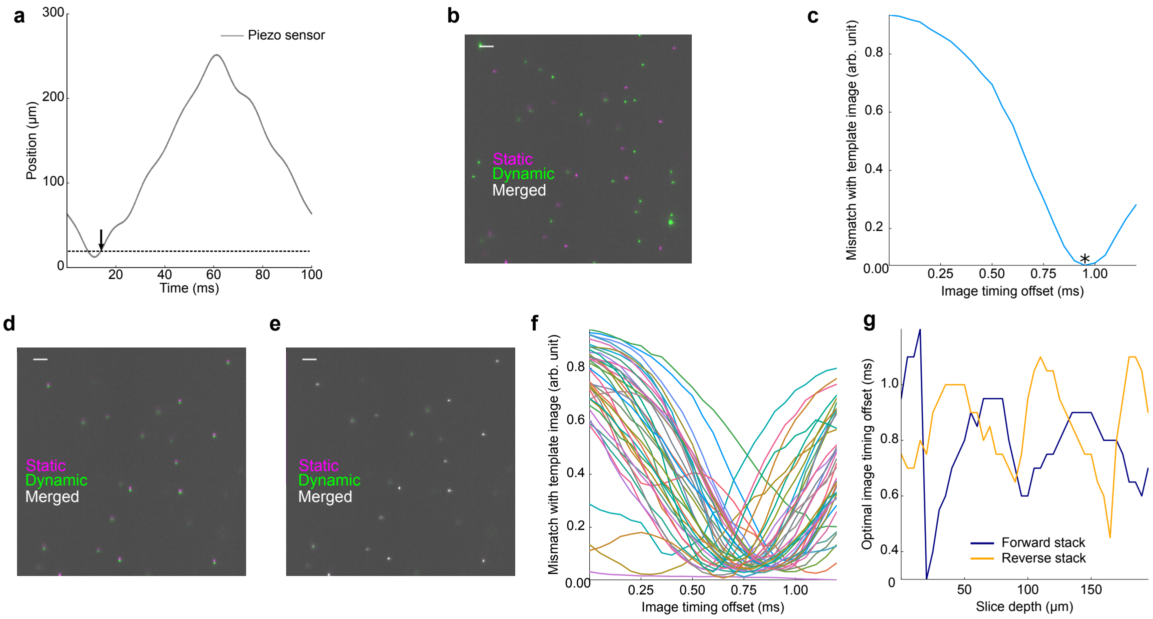


### Fig. S2

Slice timing calibration

**(a)** A single piezo response cycle as measured by the piezo's built-in capacitive sensor. The dashed line marks a position at which the user desires to acquire and image. An arrow marks the intersection of the line with the sensor trace. The timing of this intersection sets the initial guess for the timing of image acquisition. The procedure illustrated in panels b-e is applied independently for each such plane that the user desires to image.

**(b)** The initial guess at camera timing based on the piezo sensor is inaccurate for fast dynamic recordings. Two images of the same bead sample were acquired: one during dynamic operation with the timing guess described in (a), and a second image, which serves as ground truth, was acquired at the same location during a very slow scan. The overlay of the two images shows poor correspondence. Scalebar: 5 μm.

**(c)** The initial timing guess is refined by acquiring dynamic images at various temporal offsets and choosing the offset that yields an image matching the static template. The timing offset that yielded an image with minimal dissimilarity (see methods) is marked with an asterisk.

**(d)** When the images corresponding to the optimal timing from (c) are overlaid they show good axial alignment, but the dynamic image exhibits a lateral shift.

**(e)** A lateral translation is sufficient to align the dynamic image from (d) with the template. This lateral transformation is recorded and applied to align each corresponding slice of each stack in a timeseries recording.

**(f)** When the same procedure is applied to align each 5 μm-spaced slice in a stack, slices vary in their optimal timing offset.

**(g)** Optimal timing offsets are shown for each slice in both the forward and reverse stacks of a bidirectional recording. Note that optimal timings cannot be predicted by the depth in the sample, as would be expected from a simple error in sensor gain.

Fig. S3

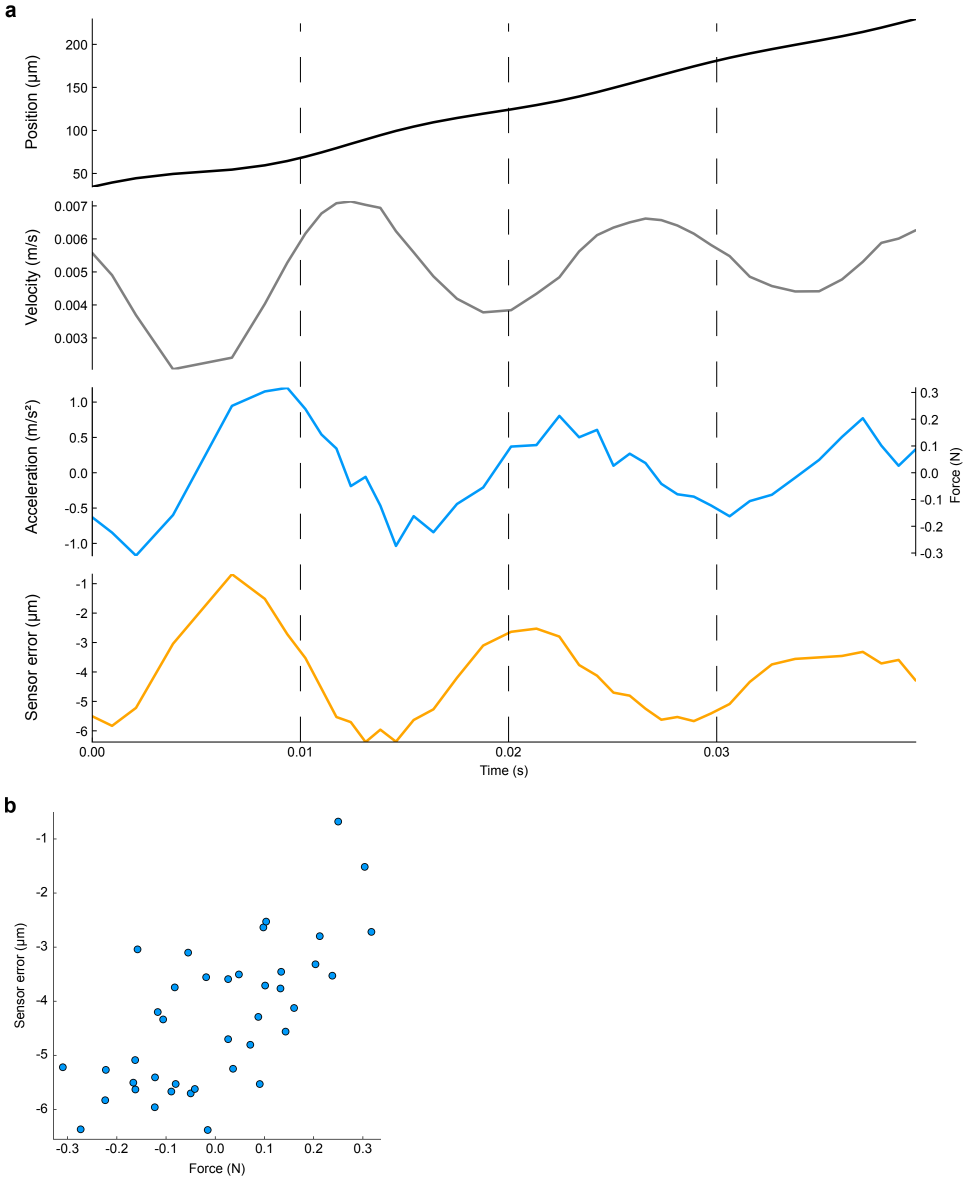


### Fig. S3

Kinetic analysis of scan system

**(a)** Error in the scanning position measurement is better predicted by the acceleration (or equivalently, force) of the scan system than by lower-order kinetics. Plotted are measured kinetic parameters of the piezo during the "forward" sweep of the cycle shown in Figure S2a. Only the subset of timepoints corresponding with the timing of each image is plotted, and the raw piezo sensor trace was first lowpass filtered with a gaussian kernel of width 300 µs to reduce sampling noise. From top to bottom: position, velocity, acceleration. The third trace also shows estimated force based only on acceleration and the 264 g load of the piezo. The fourth and final trace shows for each image slice the difference between the measured position and the actual position of the focal plane as determined with the camera in the procedure described in Figure S2. Note that this trace aligns better with the acceleration/force trace than with the other traces.

**(b)** Scatter plot of points from the blue force trace in (a) showing that force is predictive of axial displacement of the focal plane.

Fig. S4

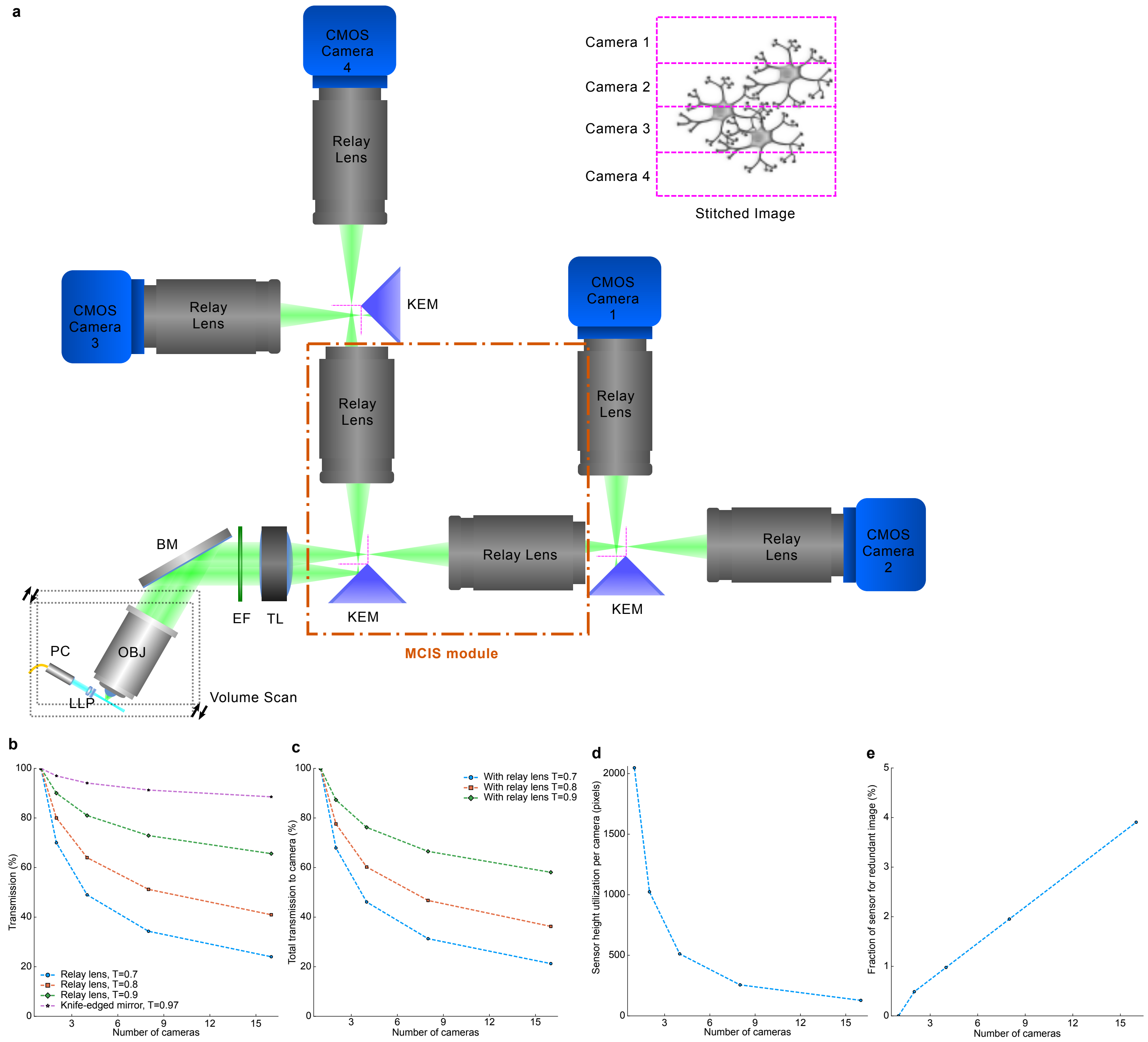


### Fig. S4

Scalability of MCIS

**(a)** Multiple MCIS modules can be chained together to split the image to more than two cameras. Shown is a diagram of an OCPI system with MCIS scaled to four cameras. The fundamental repeating unit of the design is outlined in orange. Abbreviations are the same as in Figures 1a and 3a.

**(b)** Each MCIS module compounds losses to imperfections in the transmission and reflectivity of the relay lenses and the KEM, respectively. These losses are plotted separately as a function of the number of cameras in the system for relay lens transmission efficiencies ranging from 70% to 90%.

**(c)** Same as (b) but KEM losses have been combined with transmission losses for each lens plotted.

**(d)** As more cameras are added to the system less of the vertical extent of each sensor is utilized to acquire a full stitched image. As shown in Figure 1, the maximum framerate is inversely proportional the utilized vertical extent of the sensor.

**(e)** The fraction of each image that contains redundant image information increases linearly with the number of cameras in the system. Thus the effective size of the field of view is reduced by a small amount when compared with a single-camera system.

**Fig. S5**

**a**

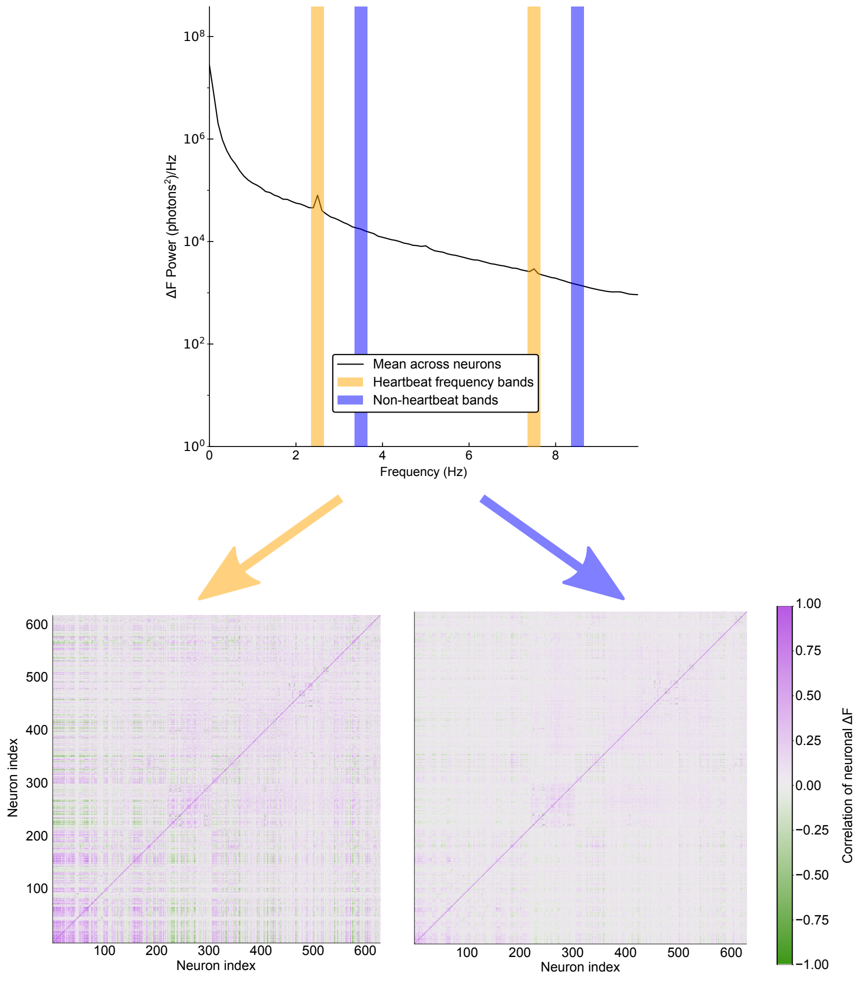
ΔF Power (photons²)/Hz
Frequency (Hz)
Mean across neurons
Heartbeat frequency bands
Non-heartbeat bands
Neuron index
Neuron index
Correlation of neuronal ΔF

**b**

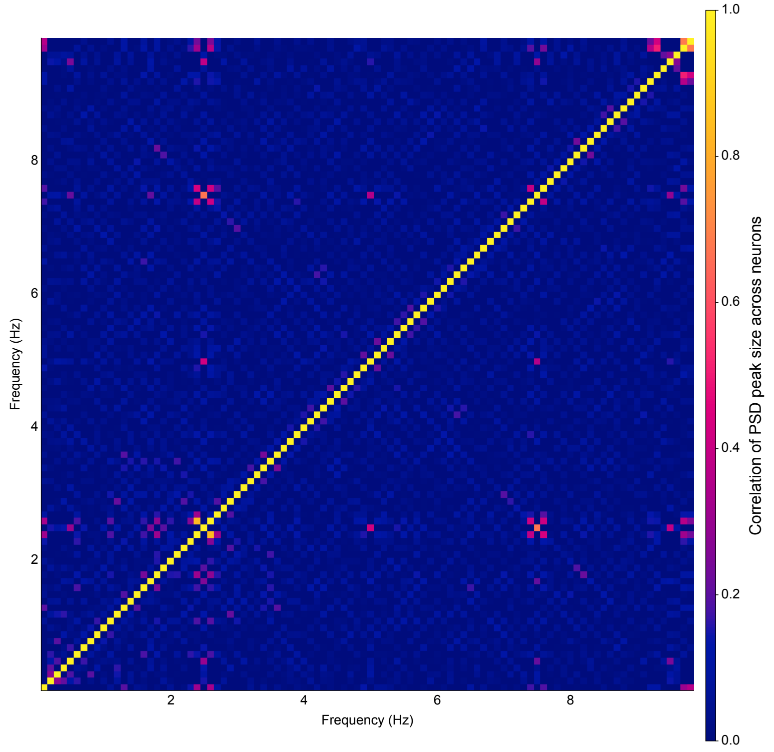
Frequency (Hz)
Frequency (Hz)
Correlation of PSD peak size across neurons

### Fig. S5

Heartbeat-mediated correlations in the ΔF signal
**(a)** Comparison of time-domain correlations in the isolated heartbeat frequency bands (left) and another nearby set of frequency bands. The former correlation matrix contains a much higher proportion of strong positive and negative correlations, as would be expected with a broadly distributed heartbeat-induced motion signal.
**(b)** The sizes of spectral deviations at the two frequencies of the putative heartbeat signal are highly correlated across neurons, suggesting that the two frequencies are components of a single signal. For each of the 629 segmented neurons we measured the size of the peak in the PSD at each frequency. Peak amplitiude was measured by dividing the PSD amplitude at each frequency by the mean amplitude of the two surrounding frequency bins (thus peak size is not defined for the maximum and minimum frequency bins). Plotted is the correlation between peak sizes across frequencies. Correlations near the diagonal are expected to be strong because they are between similar frequencies. However the strong correlation between the 7.5 Hz and 2.5 Hz peak size stands out.
