## Supplementary material for "Fast Objective Coupled Planar Illumination Microscopy": Design diagrams

**Parts List** (ordered approximately from excitation to collection of emission)

| Number in diagram | Part description | Vendor | Custom? | Part number | Price (USD) | Count | |
|---|---|---|---|---|---|---|---|
| not shown | Laser system w/5 wavelengths, AOTF | Spectral (now Andor) | no | Laser Merge Module (LMM) | 90000 | 1 | *discontinued, estimated price |
| 1 | Pigtailed fiber collimator | OZ Optics | no | LPC-01-405/640-3/125-P-1-6AC-40-3S-3-2 | 146.5 | 1 | |
| 2 | Pigtailed fiber collimator holder | (machine shop) | yes | | NA | 1 | |
| 3 | Lightsheet lens module (“cigarette”) | (machine shop) | yes | | NA | | |
| not shown | Lightsheet achromat lens | Edmund Optics | no | 45-262 | 87.5 | 1 | |
| not shown | Lightsheet cylinder lens | Tower Optical | yes* | | 99 | 1 | |
| 4 | Rotation collar | (machine shop) | | | NA | 1 | |
| 5 | Clamp for lightsheet lens holder | (machine shop) | | | NA | 1 | |
| not shown | 10x, 0.3 N.A. objective (UMPLFLN10X/W) | Olympus | no | 1-U2M583 | 774.25 | 1 | |
| 6 | 20x, 0.5 N.A. objective (UMPLFLN20X/W) | Olympus | no | 1-U2M585 | 1464.64 | 1 | |
| not shown | 40x, 0.8 N.A. objective (LUMPLFLN 40X/W) | Olympus | no | | 2500 | 1 | |
| 7 | Objective holder (RMS) | (machine shop) | yes | | | | |
| 8 | Mini-dovetail stage (2-axis) | Lightspeed Technologies Inc. | no | MDE266 | 304 | 1 | |
| 9 | Mini-dovetail stage (3-axis) | Lightspeed Technologies Inc. | no | MDE269 | 573 | 1 | |
| 10 | dovetail stabilizer | (machine shop) | yes | | NA | 1 | |
| 11 | Front piezo plate | (machine shop) | yes | | NA | 1 | |
| 12 | Piezo positioner, 800um range | Piezosystem Jena | no | NanoSX800 | 11205 | 1 | |
| not shown | Piezo amplifier (digital control) | Piezosystem Jena | no | 30DV300 | 5616 | 1 | |
| 13 | Rear piezo plate | (machine shop) | yes | | NA | | |
| not shown | Precision broadband mirror | Edmund Optics | no | 48-017 | 395 | 1 | |
| not shown | 200mm tube lens | Thorlabs | no | ITL200 | 450 | 1 | |
| not shown | Knife edged mirror | Thorlabs | no | MRAK25-G01 | 125.46 | 1 | |
| not shown | 50/50 beamsplitter (25 x 36 mm) | Thorlabs | no | BSW10R | 110 | 1 | |
| not shown | Filter cube w/insert | Thorlabs | no | DFM | 304 | 1 | |
| not shown | Extra filter cube insert | Thorlabs | no | DFMT1 | 201 | 1 | |
| not shown | 0.9x telecentric relay lens | Edmund Optics | no | 62-902 | 2265 | 2 | |
| not shown | CMOS camera | PCO | no | Edge 4.2 | 16400 | 2 | |
| not shown | Stages for aligning cameras | Thorlabs | no | DTS25 | 179.5 | 4 | |
| not shown | DAQ board (PCI-6259) | National instruments | no | 779072-01 | 1592.1 | 1 | |
| not shown | Physiology stage surface | Thorlabs | no | PHYS24BB | 2500 | 1 | |
| not shown | Lab jack | Newport | no | 281 | 999.94 | 1 | |
| not shown | Breadboard for connecting lab jack | Thorlabs | no | MB1224 | 259 | 1 | |
| not shown | XY microscope translation stage | Scientifica | no | N/A (Quote ref: QLS-30894) | 4042.5 | 1 | |
| not shown | RAID hard drives | Seagate | no | 4221403 | 78.19 | 20 | |
| not shown | Breadboard for vertical mounting of system | Thorlabs | no | MB1824 | 400 | 1 | |
| | | | | Price total: | 163760.69 | | |

***Cylindrical lens specifications**

| | |
|---|---|
| Diameter | 3mm +0/-0.2mm |
| Center Thickness | 1mm ±0.1mm |
| Edge Thickness | 1.37mm |
| Effective Focal Length | -6.25mm |
| Back Focal Length | -6.91mm |
| Focal Length Tol | ±3% |
| Radius | -3.24mm |
| Surface quality | 60/40 both sides |
| Coating | MgF2 |

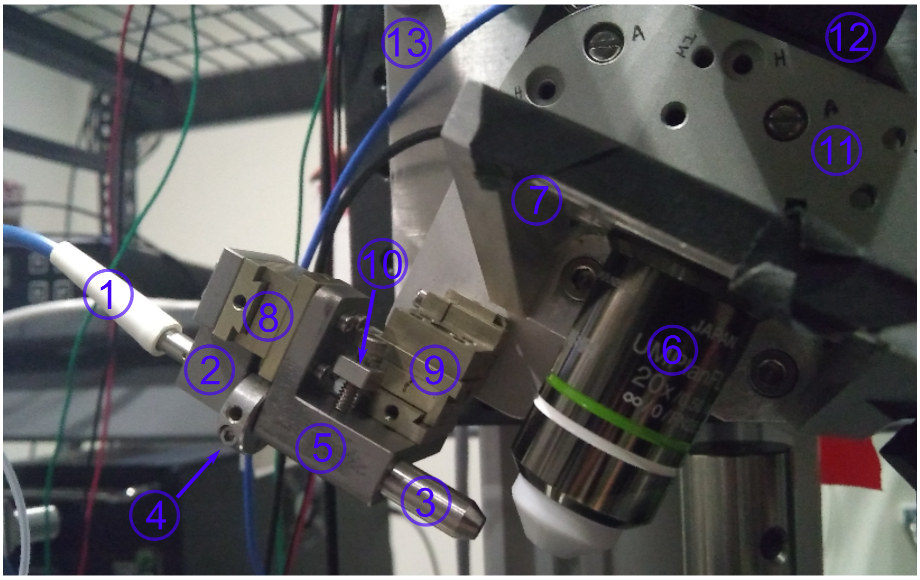


view from front

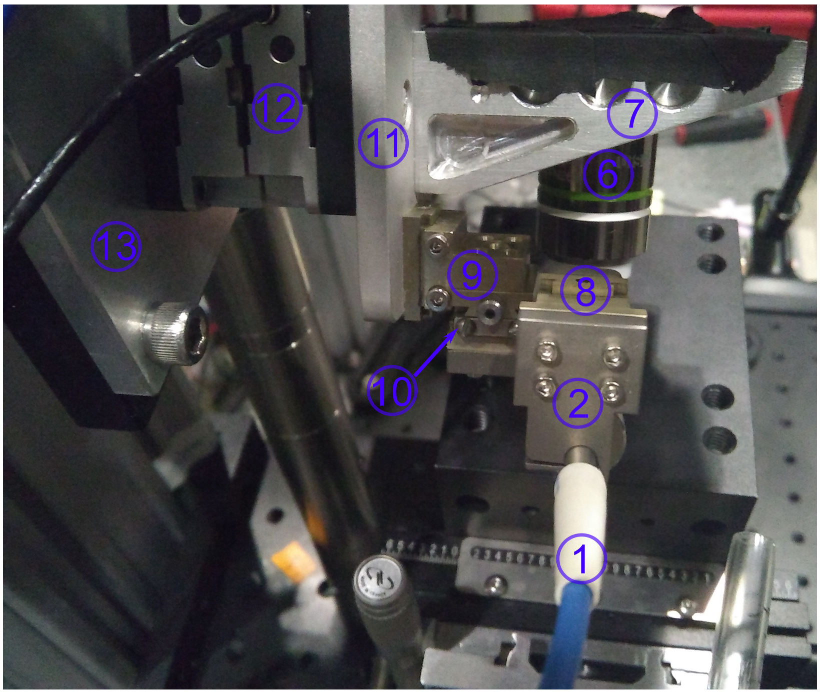


view from left

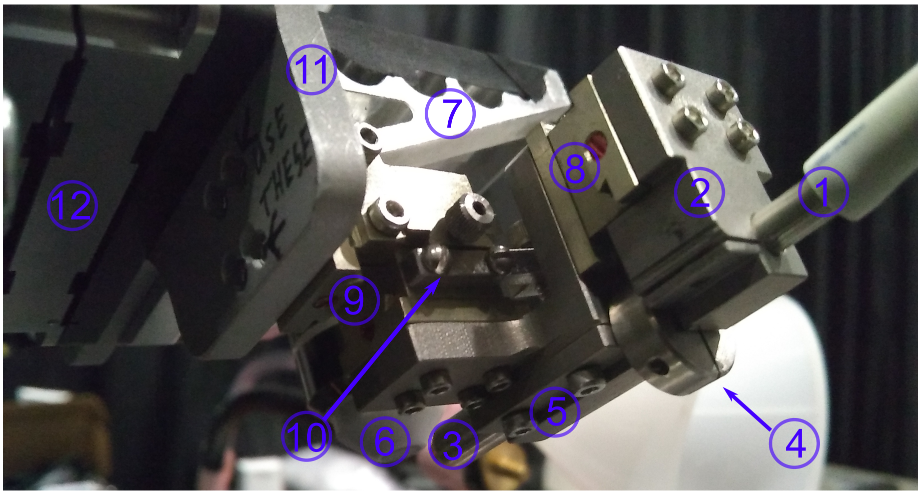


view from rear left

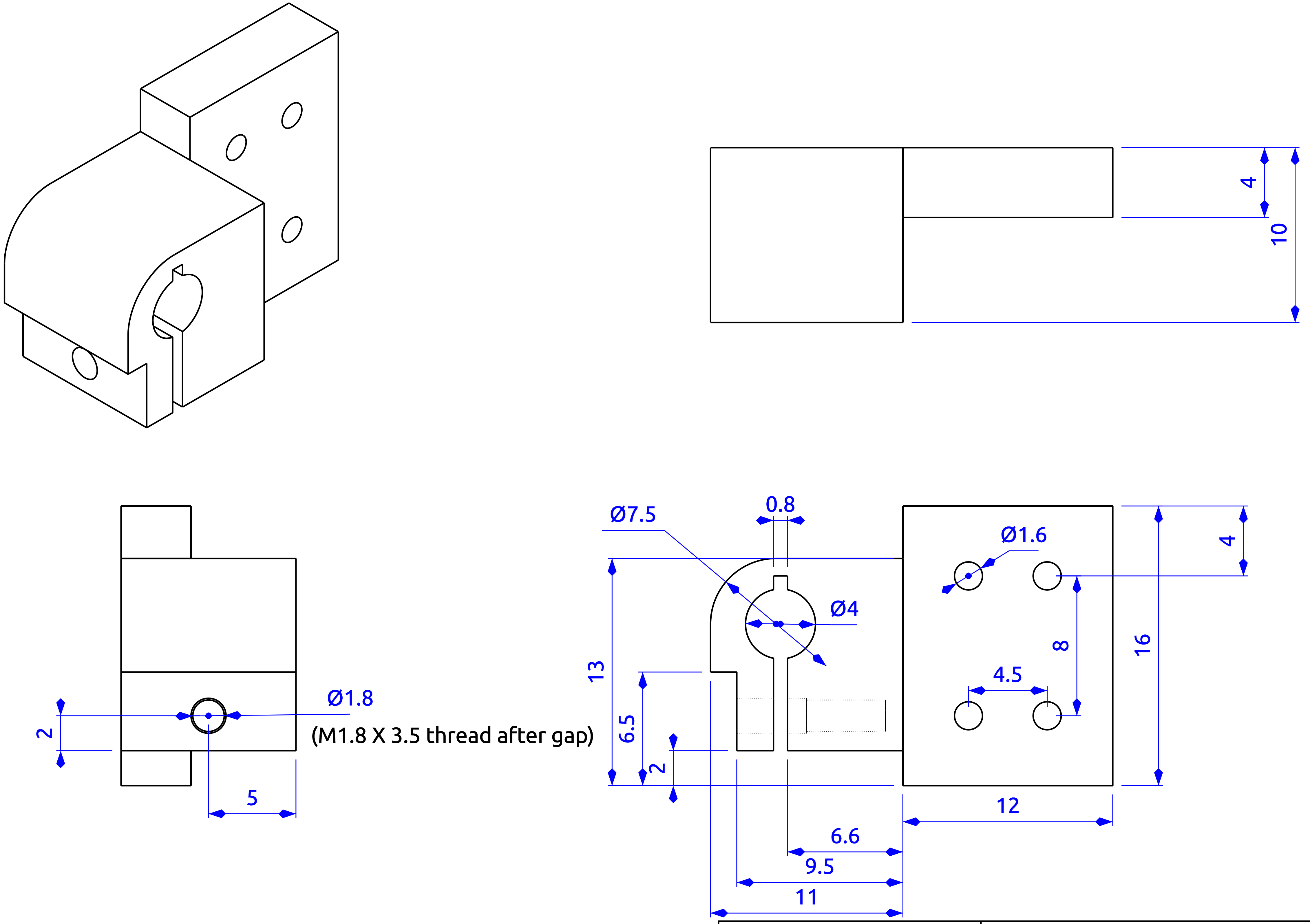


| Created by: | Title: | | |
|---|---|---|---|
| Cody Greer | Pigtailed collimator holder | | |
| Supplementary information: | Size: A3 | Sheet: X / Y | Scale: mm |
| Clamps onto pigtailed collimator<br>Attaches to MDE266 dovetail slide<br>Addtl authors: T.E. Holy J. Kreitler K. Poenicke | Part number: NA | | |
| | Drawing number: DN | | |
| | Date: 17/10/2017 | | Revision: REV A |

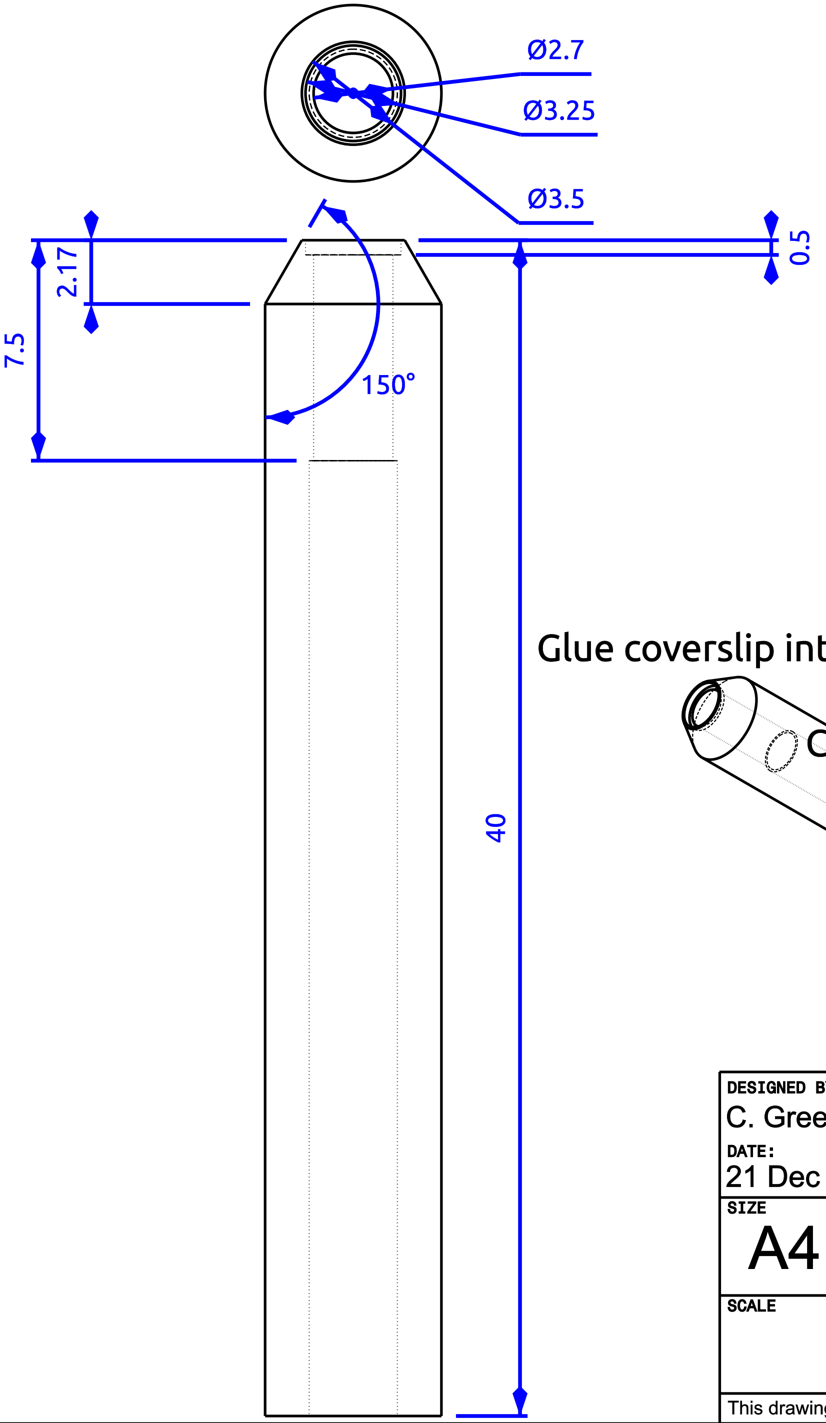


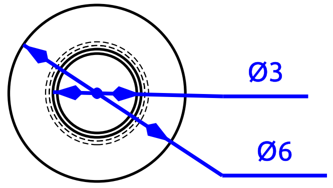


Glue coverslip into recession at tip

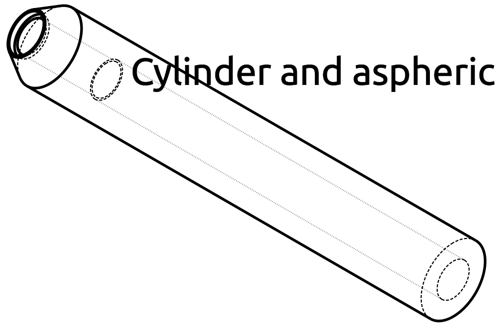

Cylinder and aspheric lenses rest on this ledge

Insert lenses from this side

| | | | |
|---|---|---|---|
| DESIGNED BY: C. Greer, T.E. Holy<br>DATE: 21 Dec 2017 | | Lightsheet Lens Module<br>"cigarette" | |
| SIZE<br>A4 | | | |
| SCALE | WEIGHT (kg) | DRAWING NUMBER<br>Addtl designers: J. Kreitler, K. Poenicke | SHEET |

This drawing is our property; it can't be reproduced or communicated without our written consent.

| | |
|---|---|
| G | _ |
| F | _ |
| E | _ |
| D | _ |
| C | _ |
| B | _ |
| A | _ |

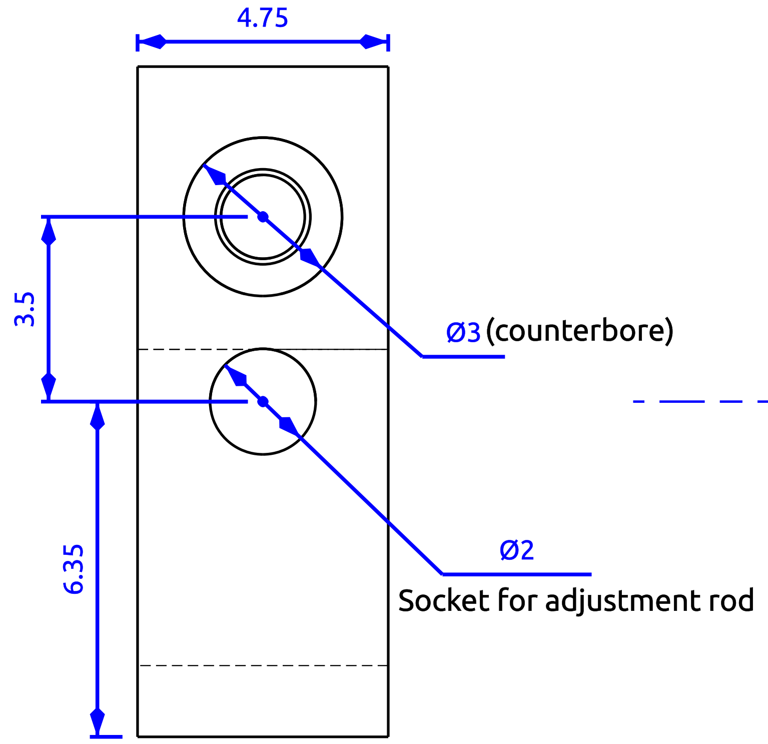


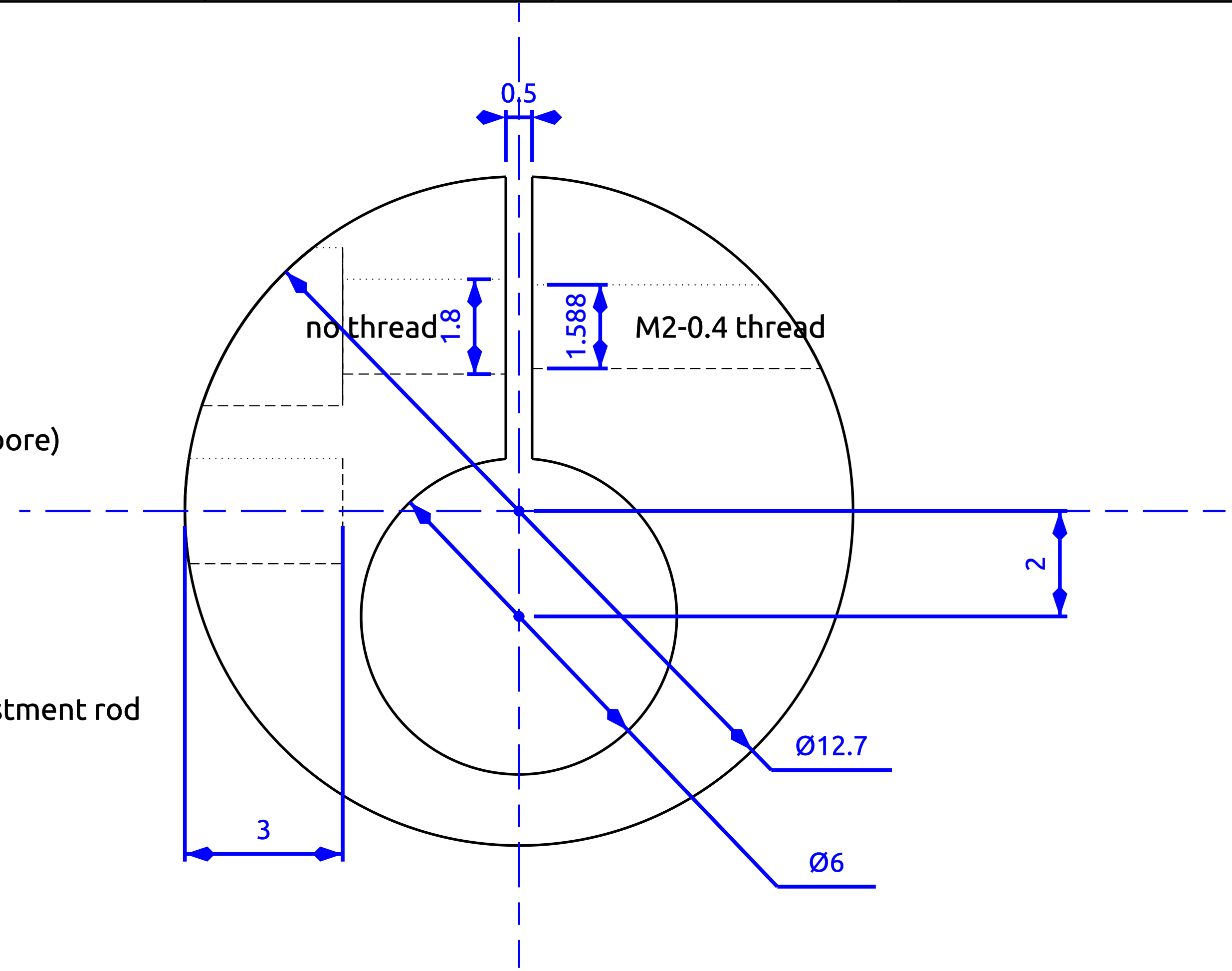


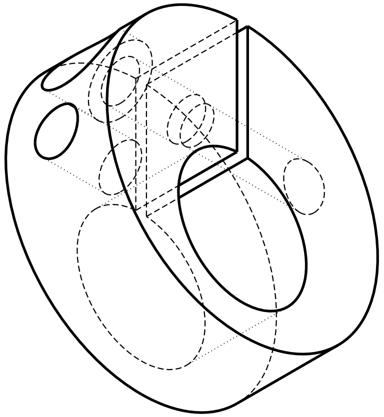

**DESIGNED BY:**
C. Greer, T. E. Holy

**DATE:**
22 December 2017

### Rotation Collar

Attaches to lightsheet "cigarette"

**SIZE**
A4

**SCALE**

**WEIGHT (kg)**

**DRAWING NUMBER**
Addtl designers: J. Kreitler, K. Poenicke

**SHEET**

| | |
|---|---|
| G | _ |
| F | _ |
| E | _ |
| D | _ |
| C | _ |
| B | _ |
| A | _ |

This drawing is our property; it can't be reproduced or communicated without our written consent.

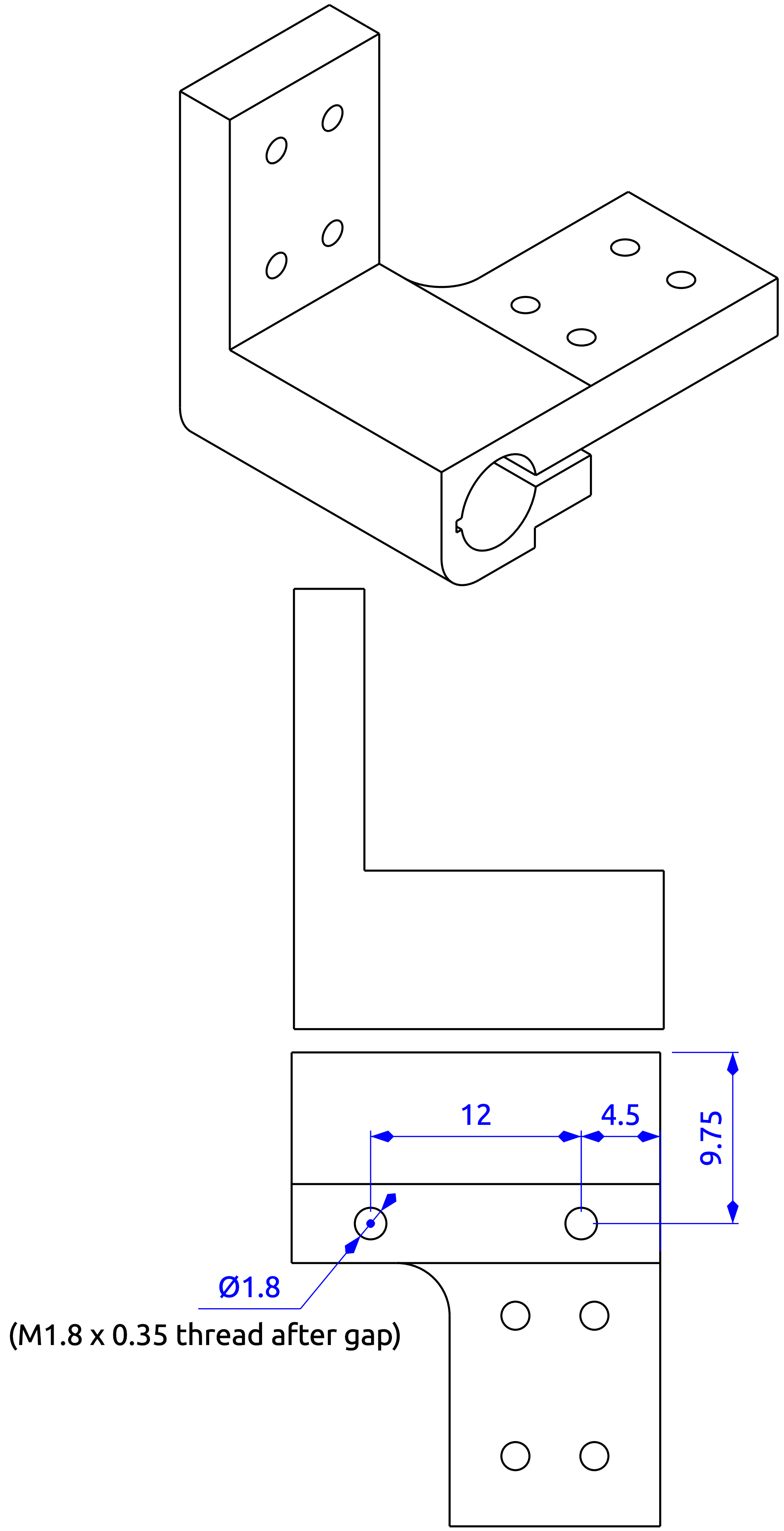


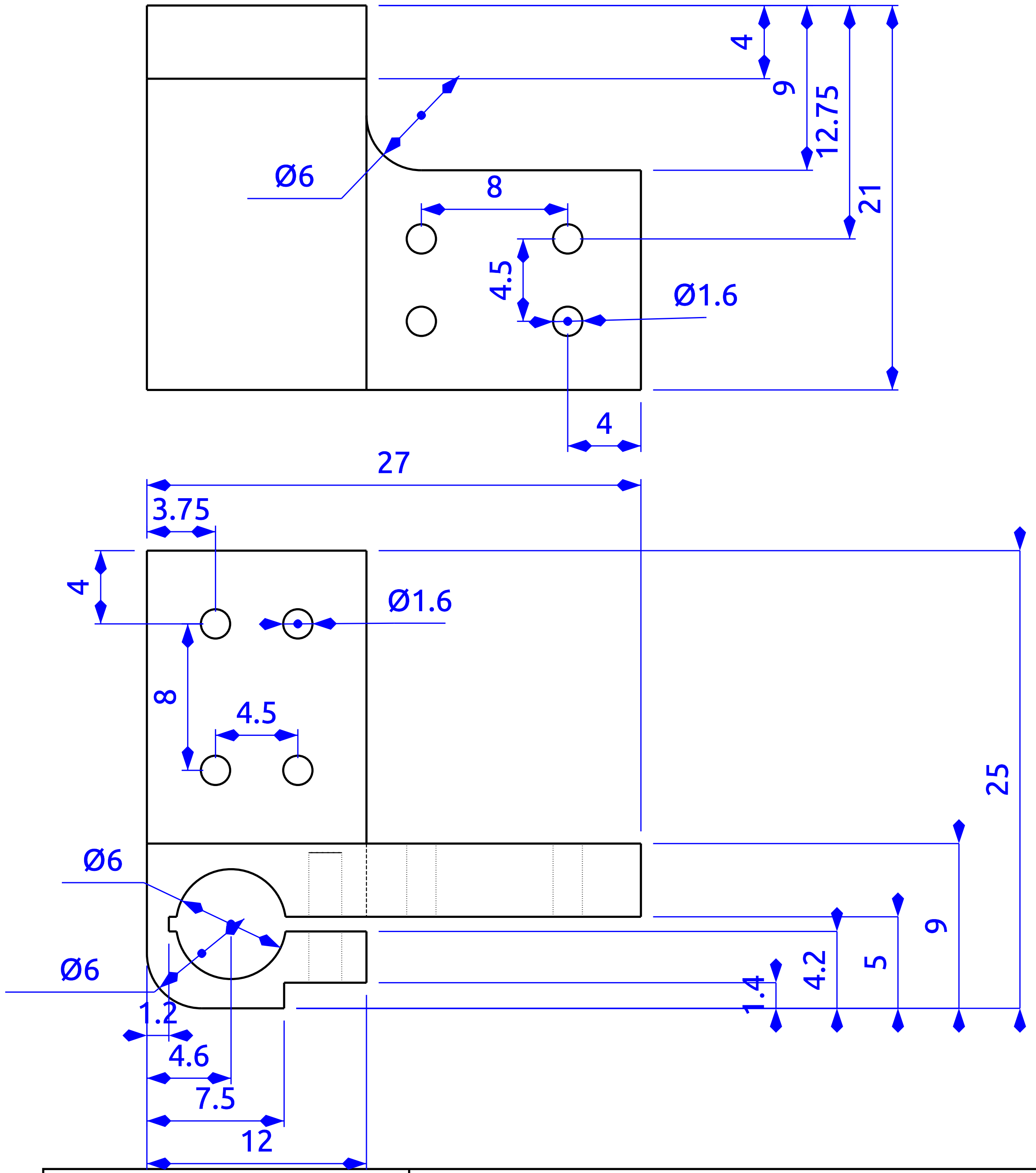


| | | | |
|---|---|---|---|
| Created by: Cody Greer | Title: Clamp for light-sheet lens holder | | |
| Supplementary information: Clamps around the light-sheet "cigarette" attaches to dovetail slides MDE266 MDE269 Addtl authors: T. E. Holy J. Kreitler K. Poenicke | Size: A3 | Sheet: X / Y | Scale: mm |
| | Part number: | | |
| | Drawing number: NA | | |
| | Date: 18/10/2017 | | Revision: REV A |

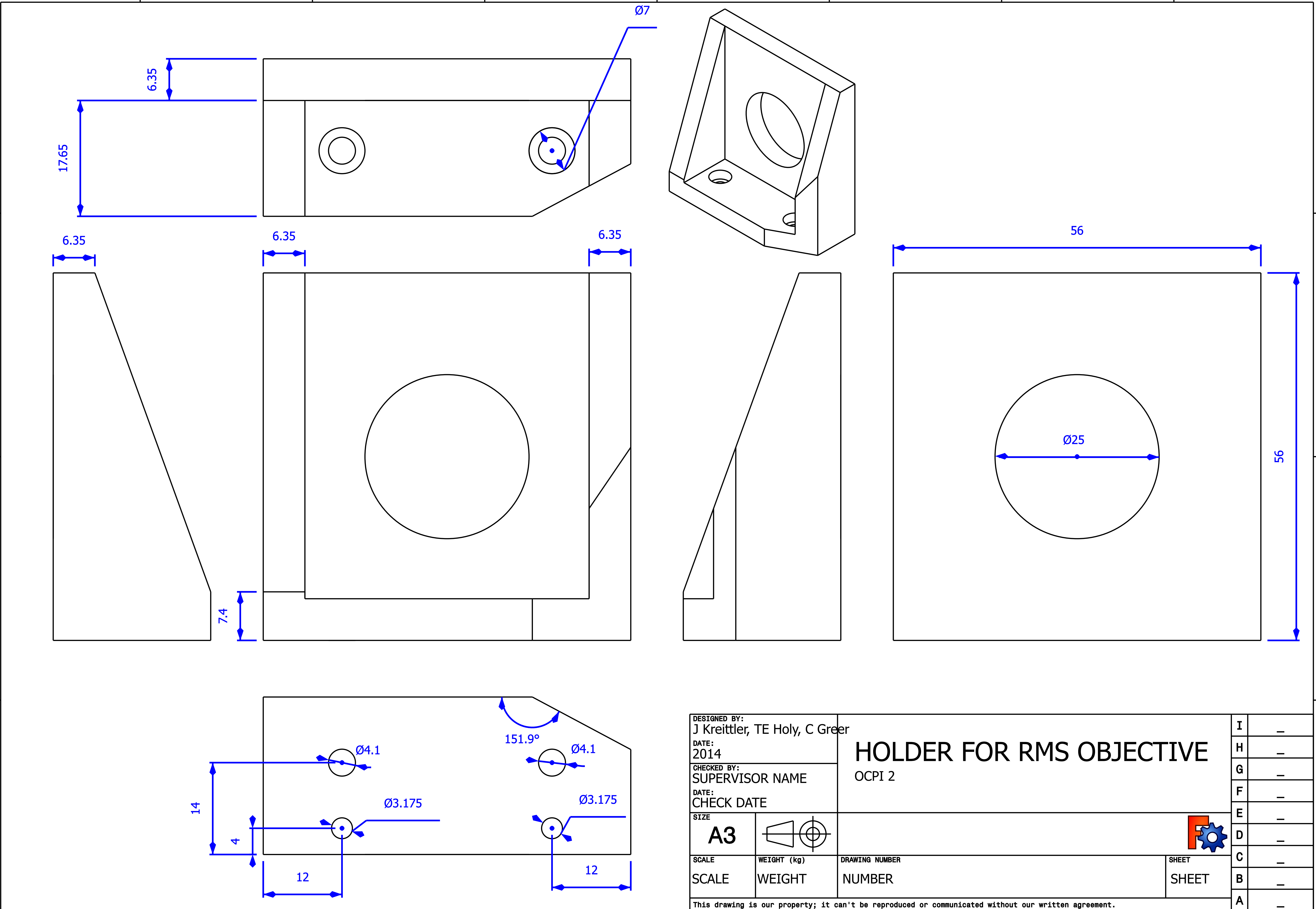


DESIGNED BY:
J Kreittler, TE Holy, C Greer
DATE:
2014
CHECKED BY:
SUPERVISOR NAME
DATE:
CHECK DATE

### HOLDER FOR RMS OBJECTIVE

OCPI 2

SIZE: A3

| SCALE | WEIGHT (kg) | DRAWING NUMBER | SHEET |
|---|---|---|---|
| SCALE | WEIGHT | NUMBER | SHEET |

This drawing is our property; it can't be reproduced or communicated without our written agreement.

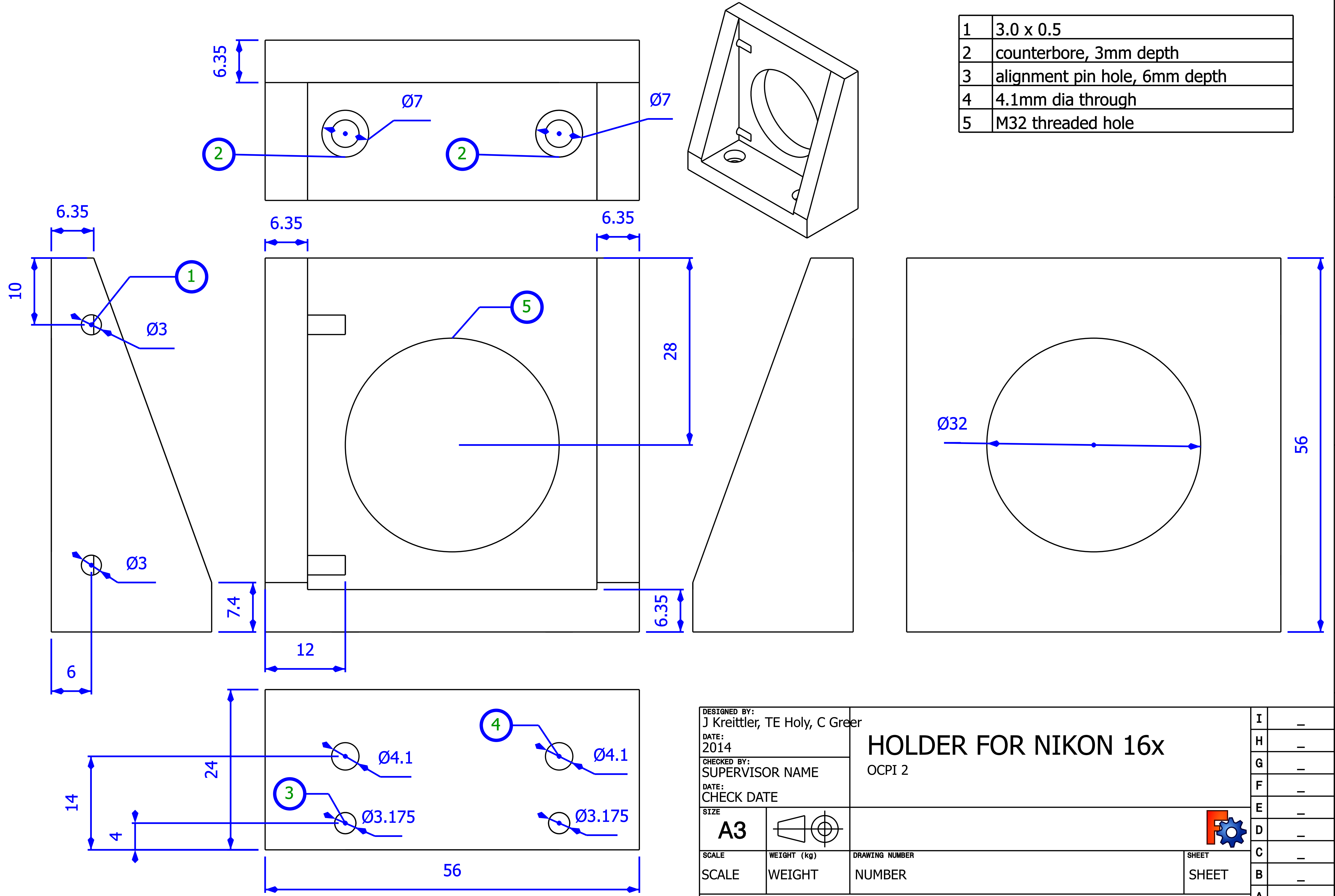


| | |
|---|---|
| 1 | 3.0 x 0.5 |
| 2 | counterbore, 3mm depth |
| 3 | alignment pin hole, 6mm depth |
| 4 | 4.1mm dia through |
| 5 | M32 threaded hole |

DESIGNED BY:
J Kreittler, TE Holy, C Greer
DATE:
2014
CHECKED BY:
SUPERVISOR NAME
DATE:
CHECK DATE

### HOLDER FOR NIKON 16x

OCPI 2

SIZE: A3

SCALE: SCALE
WEIGHT (kg): WEIGHT
DRAWING NUMBER: NUMBER
SHEET: SHEET

This drawing is our property; it can't be reproduced or communicated without our written agreement.

| | |
|---|---|
| 1 | 3.0 x 0.5 |
| 2 | counterbore, 3mm depth |
| 3 | alignment pin hole, 6mm depth |
| 4 | 4.1mm dia through |
| 5 | M34 threaded hole |

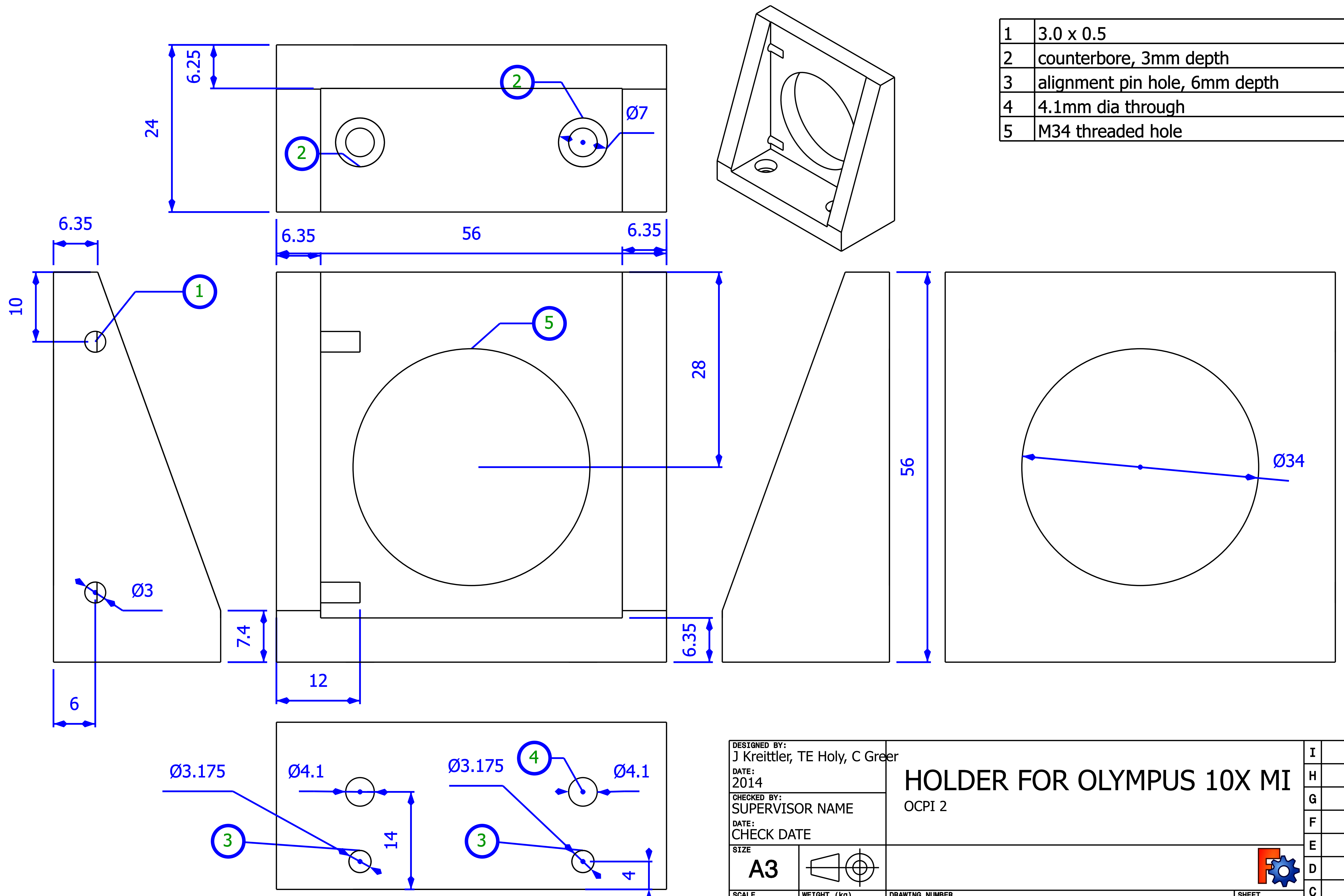


DESIGNED BY:
J Kreittler, TE Holy, C Greer
DATE:
2014
CHECKED BY:
SUPERVISOR NAME
DATE:
CHECK DATE

### HOLDER FOR OLYMPUS 10X MI

OCPI 2

| SIZE | SCALE | WEIGHT (kg) | DRAWING NUMBER | SHEET |
|---|---|---|---|---|
| A3 | SCALE | WEIGHT | NUMBER | SHEET |

This drawing is our property; it can't be reproduced or communicated without our written agreement.

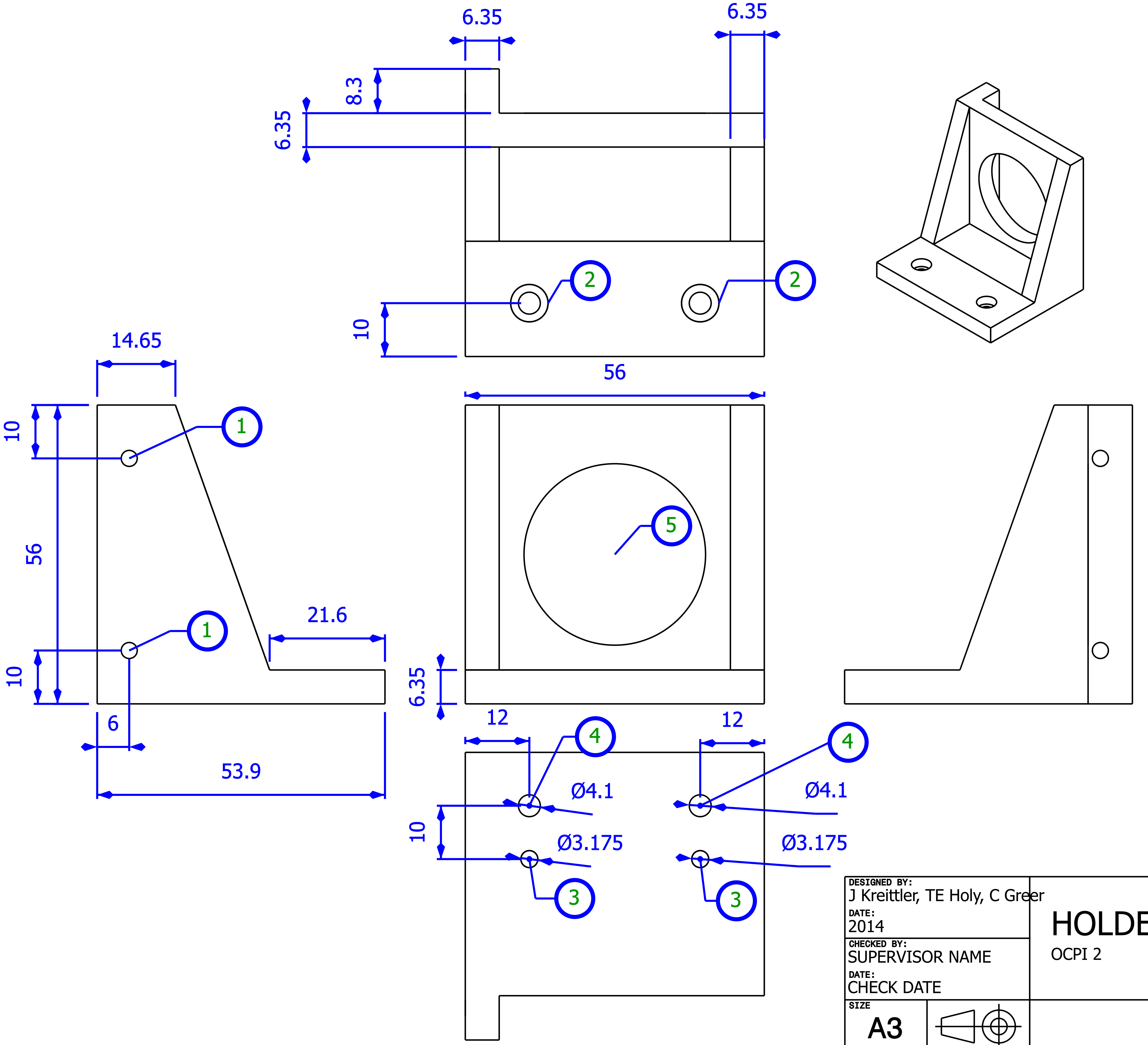


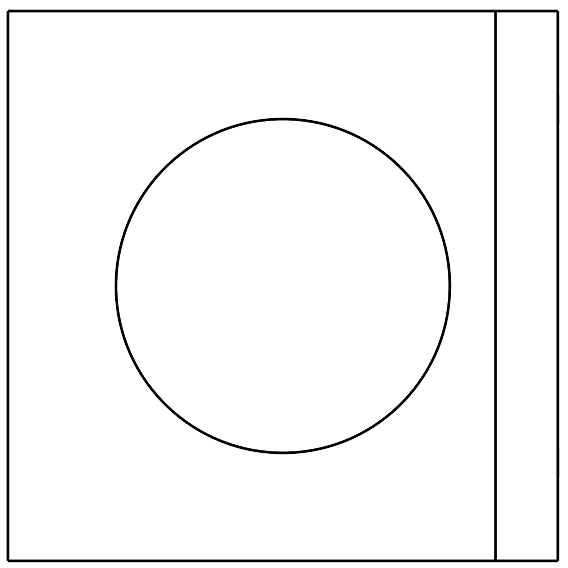

| | |
|---|---|
| 1 | 3.0 x 0.5 thread |
| 2 | 7mm dia counterbore, 3mm depth |
| 3 | alignment pin hole, 6mm depth |
| 4 | 4.1mm dia through |
| 5 | M34 threaded hole for objective |

DESIGNED BY:
J Kreittler, TE Holy, C Greer
DATE:
2014
CHECKED BY:
SUPERVISOR NAME
DATE:
CHECK DATE

### HOLDER FOR OLYMPUS 4x

OCPI 2

SIZE: A3

SCALE: SCALE | WEIGHT (kg): WEIGHT | DRAWING NUMBER: NUMBER | SHEET: SHEET

| | |
|---|---|
| I | _ |
| H | _ |
| G | _ |
| F | _ |
| E | _ |
| D | _ |
| C | _ |
| B | _ |
| A | _ |

This drawing is our property; it can't be reproduced or communicated without our written agreement.

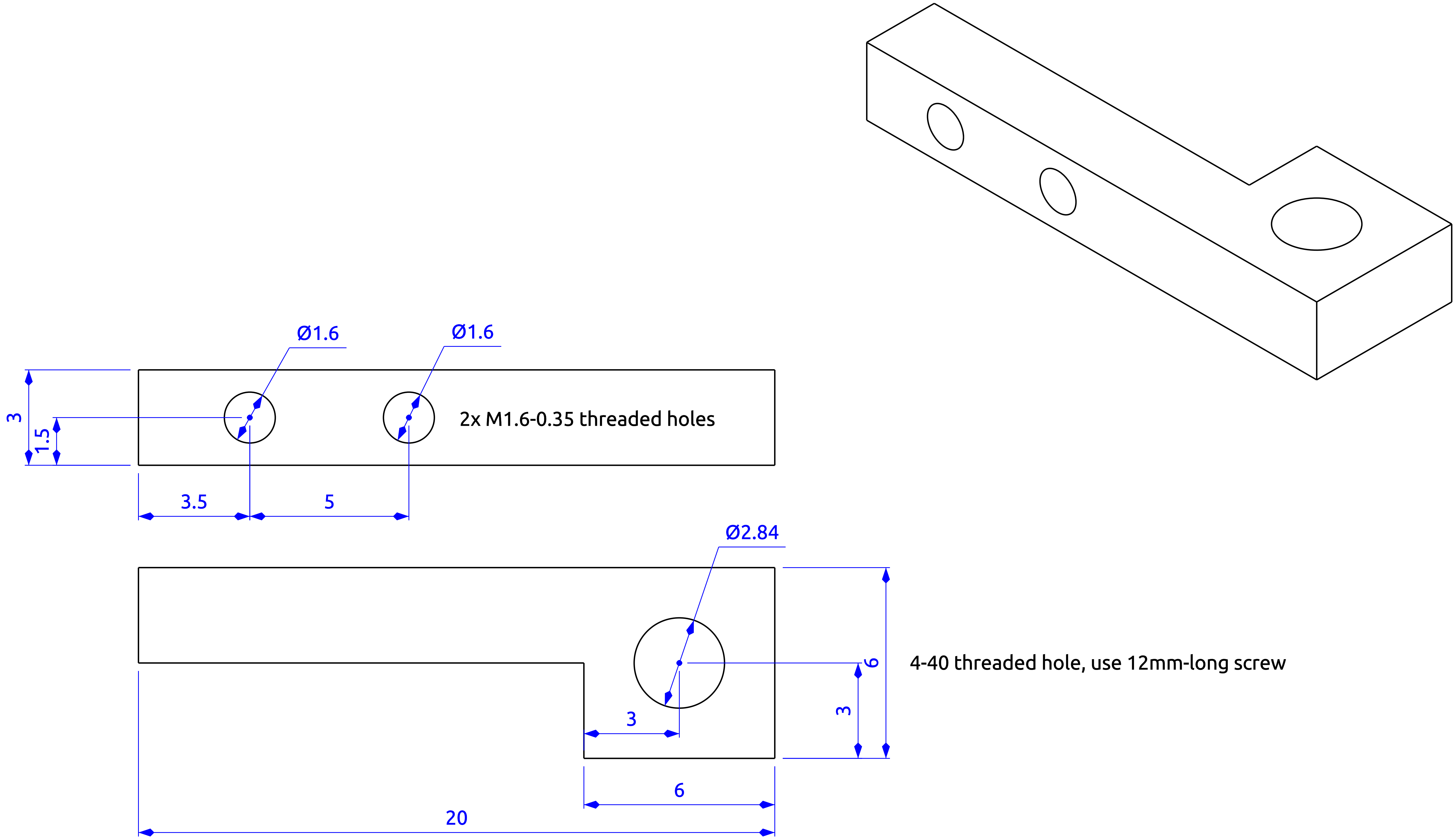


DESIGNED BY:
C. Greer, K. Poenicke
DATE:
1 November 2017
CHECKED BY:
NA
DATE:
NA

### Dovetail stabilizer

Applies tension to dovetail joint

SIZE
A3

| SCALE | WEIGHT (kg) | DRAWING NUMBER | SHEET |
| --- | --- | --- | --- |
| mm | NA | NA | NA |

This drawing is our property; it can't be reproduced or communicated without our written agreement.

| | |
| --- | --- |
| I | _ |
| H | _ |
| G | _ |
| F | _ |
| E | _ |
| D | _ |
| C | _ |
| B | _ |
| A | _ |

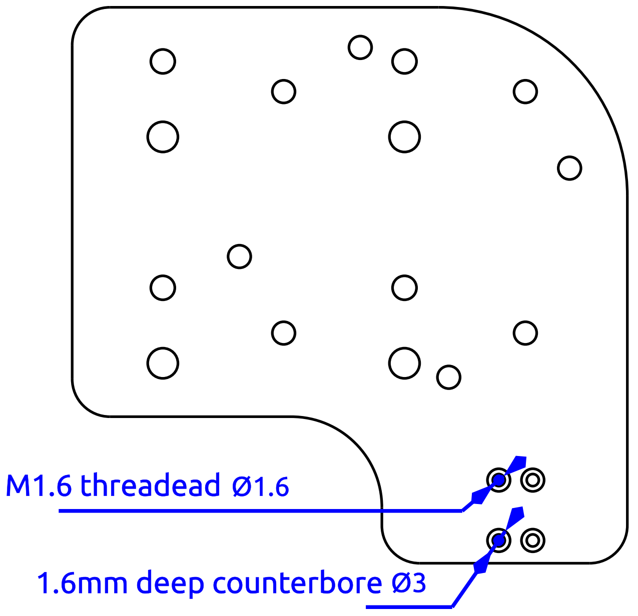


Rear view

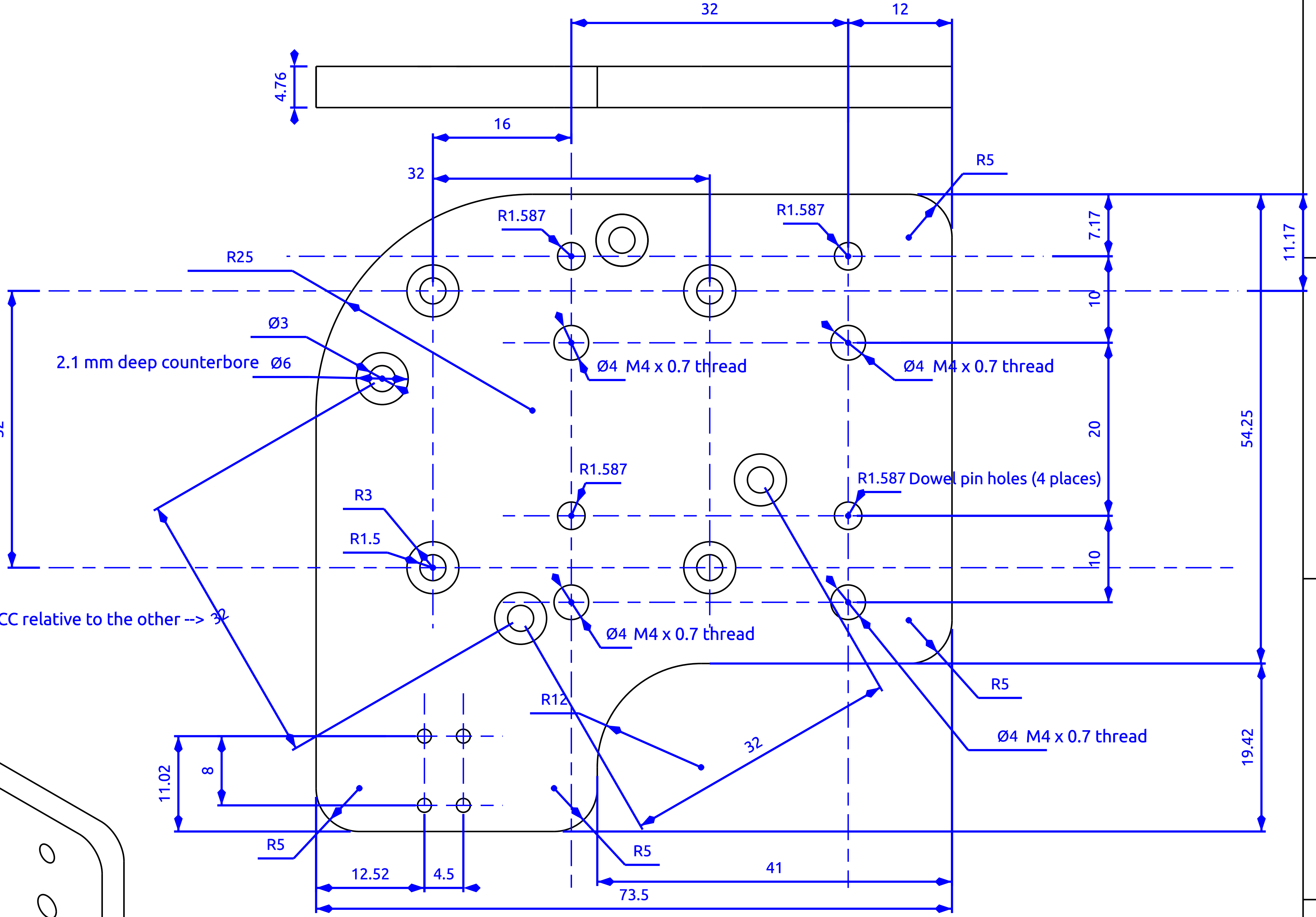


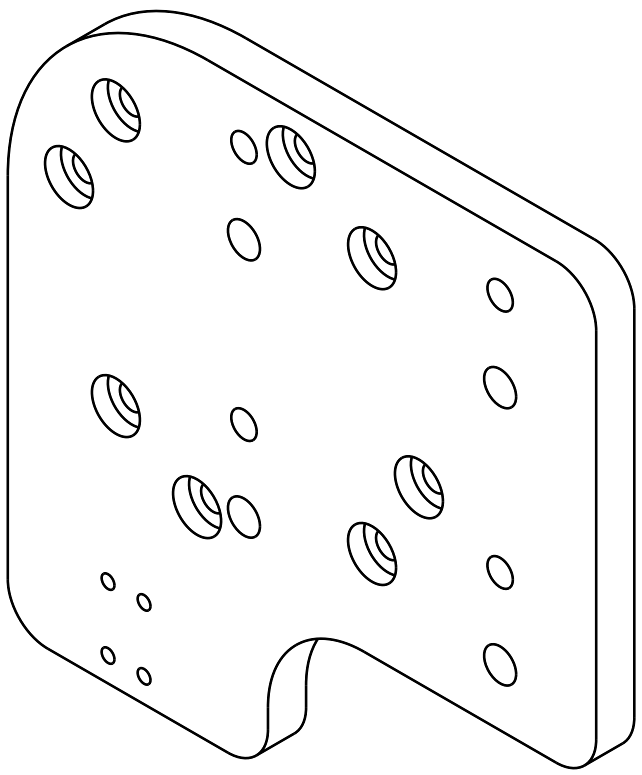

DESIGNED BY:
Cody Greer, John Kreittler, Kevin Poenicke

DATE:
19 February 2017

CHECKED BY:
SUPERVISOR NAME

DATE:
CHECK DATE

### Front piezo plate

OCPI 2 microscope

SIZE: A3

SCALE: SCALE

WEIGHT (kg): WEIGHT

DRAWING NUMBER: NUMBER

SHEET: SHEET

| | |
|---|---|
| I | _ |
| H | _ |
| G | _ |
| F | _ |
| E | _ |
| D | _ |
| C | _ |
| B | _ |
| A | _ |

This drawing is our property; it can't be reproduced or communicated without our written agreement.

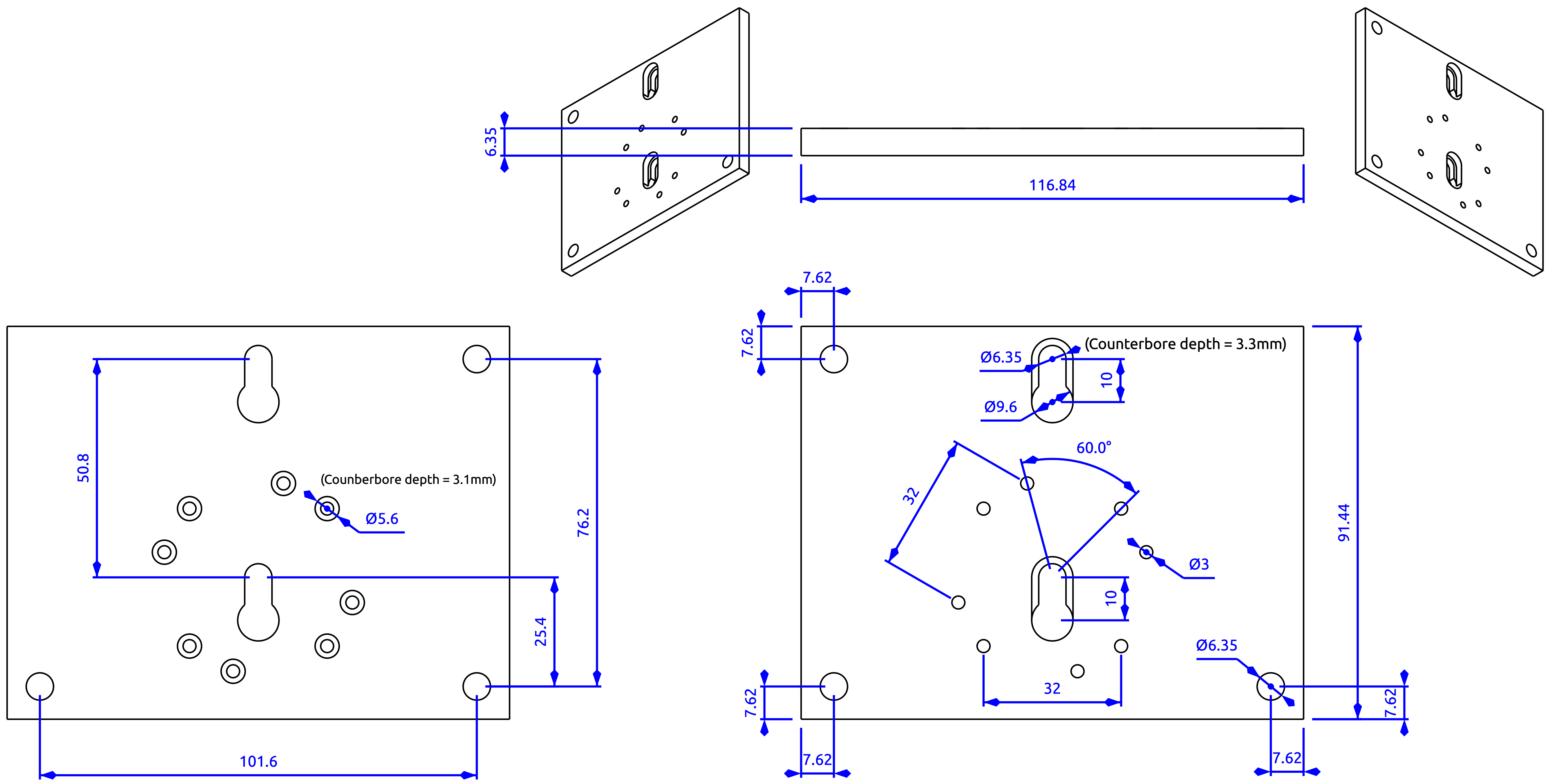


| | | | |
|---|---|---|---|
| DESIGNED BY: Cody Greer | Rear Piezo Plate | | |
| DATE: 19 February 2017 | OCPI 2 Microscope | | |
| CHECKED BY: SUPERVISOR NAME | | | |
| DATE: CHECK DATE | | | |
| SIZE A3 | | | |
| SCALE SCALE | WEIGHT (kg) WEIGHT | DRAWING NUMBER NUMBER | SHEET SHEET |

This drawing is our property; it can't be reproduced or communicated without our written agreement.
